# UV photogrammetry for transparent and composite surfaces

**DOI:** 10.64898/2026.04.20.719583

**Authors:** Gregor J. Gentsch, Adrian Platz, Matthew Guo, Lukas Harder, Dennis Boettger, Gunnar Brehm, Christian Franke, Andreas W. Stark

**Author notes:** These authors contributed equally to this work and have the right to list their name first in their CV.

## Abstract

Transparent and composite surfaces pose a fundamental challenge for stereo photogrammetry: optically smooth glass produces no detectable surface features under visible illumination, making three-dimensional reconstruction impossible without surface preparation. This excludes optical components such as lenses and cover glasses, composite assemblies, and semi-translucent biological specimens from non-contact geometric measurement. Here we show that coherent speckle illumination at 266 nm overcomes this limitation by exploiting wavelength-dependent scatter enhancement, generating sufficient backscattered signal on surfaces that are entirely invisible under visible illumination. We developed a multispectral stereo system and evaluated three illumination modalities under identical acquisition conditions. On transparent glass, both visible modalities produce complete reconstruction failure, recovering only non-transparent holder structures. Ultraviolet speckle illumination at 266 nm enables dense reconstruction of the same surfaces. We demonstrate recovery of an uncoated plano-convex lens with a fitted radius of 30.946 mm and point-cloud standard deviation of 106.5 µm, defect detection on a transparent cover glass without surface preparation, and reconstruction of a semi-translucent biological specimen. On metrology-grade reference objects, ultraviolet speckle achieves a standard deviation of 116 µm and completeness exceeding 93%, approaching the performance of optimised visible structured illumination. These results establish ultraviolet speckle photogrammetry as an enabling approach of optical metrology to otherwise uncooperative surfaces, with relevance to optical manufacturing inspection and biological surface analysis.

## Introduction

Optical 3D surface metrology with active illumination plays a central role in modern manufacturing, quality control, and biological imaging^1^. Photogrammetry and stereo reconstruction are particularly attractive because they allow flexible measurements of complex geometries using comparatively simple hardware. Techniques such as fringe projection^1^, aperiodic fringe projection^2^, and speckle projection^3–5^ are widely used because they enable rapid, non-contact measurements of complex surfaces, even for biological specimens, including three-dimensional reconstruction of mouse vibrissal arrays used to geometrically verify cortical topography^6,7^. However, these methods are based on the reliable detection of corresponding image features in multiple views and thus their performance strongly depends on the optical properties of the measured object. Transparent or weakly scattering surfaces typically lack such features under VIS illumination, preventing robust correspondence estimation^8,9^. A particularly challenging class of objects are surfaces that simultaneously exhibit transparent, reflective, and scattering regions. In the following, we refer to such objects as composite surfaces. Typical examples include optical components such as lenses and cover glasses, biological specimens with transparent structures, or technical objects combining glass and rough materials.

For only reflective surfaces, deflectometry^10–13^ has become a metrological tool to maintain the same advantages of contact-lessness and speed while achieving even higher precision than scattering based photometry. On the other hand, it becomes unreliable when strong scattering occurs. Interferometric techniques^14,15^ can achieve extremely high precision but typically require well-controlled optical conditions and are difficult to apply to objects with mixed optical properties or large geometrical variations. Transparent objects, such as glass samples, can be measured in their 3D shape when infrared illumination is used and the self-emitted thermal radiation from the object surface is imaged for 3D triangulation. This technique can also be used to measure some composite objects ^16,17^. In comparison to scattering based techniques this approach utilizes the self-emitted radiation from the objects, making it largely independent of optical scattering properties and therefore well suited for transparent and specular materials. However, the achievable spatial resolution and temporal response of such infrared-based approaches can be limited by thermal diffusion processes and material-dependent absorption, which may reduce the visibility of fine surface features and complicate measurements on heterogeneous or weakly absorbing specimens.

One possible strategy to overcome the limitation of stereo photogrammetry in the visible range, while still relying on scattered illumination and including composite surfaces, is to exploit the wavelength dependence of optical scattering. For many materials, scattering strength increases significantly towards shorter wavelengths. For optically smooth surfaces, scatter intensity in the Beckmann-Kirchhoff regime scales approximately as the square of the ratio of surface roughness to wavelength, predicting a substantial backscatter increase towards ultraviolet wavelengths independent of bulk material properties^18^. For subsurface inhomogeneities and bulk contributions, Rayleigh-type scaling provides an additional enhancement with an approximate λ⁻⁴ dependence^18,19^. Both mechanisms act simultaneously in optical glass, suggesting that ultraviolet illumination may strongly enhance the visibility of otherwise transparent surfaces and enable stereo reconstruction where visible illumination fails entirely.

In this work, we investigated the use of UV speckle illumination to enable stereo photogrammetric reconstruction of transparent and composite surfaces. We developed a measurement system capable of operating in both the VIS and UV spectral ranges and evaluated it using three illumination modalities: incoherent broadband VIS structured patterns, coherent speckle illumination at 532 nm, and UV speckle illumination at 266 nm. The comparison of coherent and incoherent illumination in the visible range further provides initial indications on the influence of coherence properties on reconstruction performance, suggesting potential benefits of extending ultraviolet illumination towards incoherent projection schemes. The approach is validated using metrology-grade reference objects, including planes and spheres, as well as challenging real-world examples such as glass lenses, cover glasses, composite objects and partially translucent biological specimens. The experiments demonstrate that UV speckle illumination produces detectable back-scattered signals even on optically smooth transparent materials, thereby enabling reliable correspondence estimation and robust 3D reconstruction. Beyond enabling geometric reconstruction of composite surfaces, the proposed approach takes classical stereo photogrammetry towards practical inspection scenarios by facilitating defect detection and pre-screening capabilities, thereby supporting efficient 100% quality control and complementing high-precision techniques such as deflectometry or interferometry.

## Results

Transparent and composite objects are inaccessible to conventional stereo photogrammetry because VIS illumination produces no detectable surface features on optically smooth glass^8,9^. The experiments reported here use three illumination modalities: incoherent broadband VIS structured illumination, coherent speckle at 532 nm, and coherent speckle at 266 nm. The VIS projector serves as a cooperative-surface baseline, it is not optimized for transparent objects and is not the primary comparison axis. The controlled comparison is between the two coherent speckle modalities, which share identical acquisition parameters, differing only in wavelength. Our central result is unambiguous: on transparent glass surfaces, both VIS modalities, structured and 532 nm speckle, produce complete reconstruction failure. Not degraded or partial reconstruction, but no sensible recovery of the transparent surface itself; only non-transparent holder structures are detected. This categorical failure is not a matter of degree. UV speckle illumination at 266 nm eliminates it entirely, enabling dense reconstruction of the same surfaces. Under both VIS structured illumination and coherent 532 nm speckle illumination, the lens surface appears entirely featureless in the camera images, correspondence estimation fails, and reconstruction returns only sparse points associated with the non-transparent holder, but the transparent glass itself is not recovered at any point. This is not a noise or threshold effect: no signal from the lens surface is present in either VIS modality. In contrast, UV speckle illumination at 266 nm produces clear spatial intensity variations across the lens surface (Fig. 1b), enabling robust correlation-based matching and dense 3D reconstruction (Fig. 1d). Quantitative evaluation confirms the geometric accuracy of the reconstruction. A best-fit sphere yields a radius of 30.946 mm with a standard deviation of 106.5 µm (Fig. 1e, f), in good agreement with the nominal geometry. Remaining gaps near the center originate from specular reflections simultaneously observed in both camera views.

**Figure 1.**
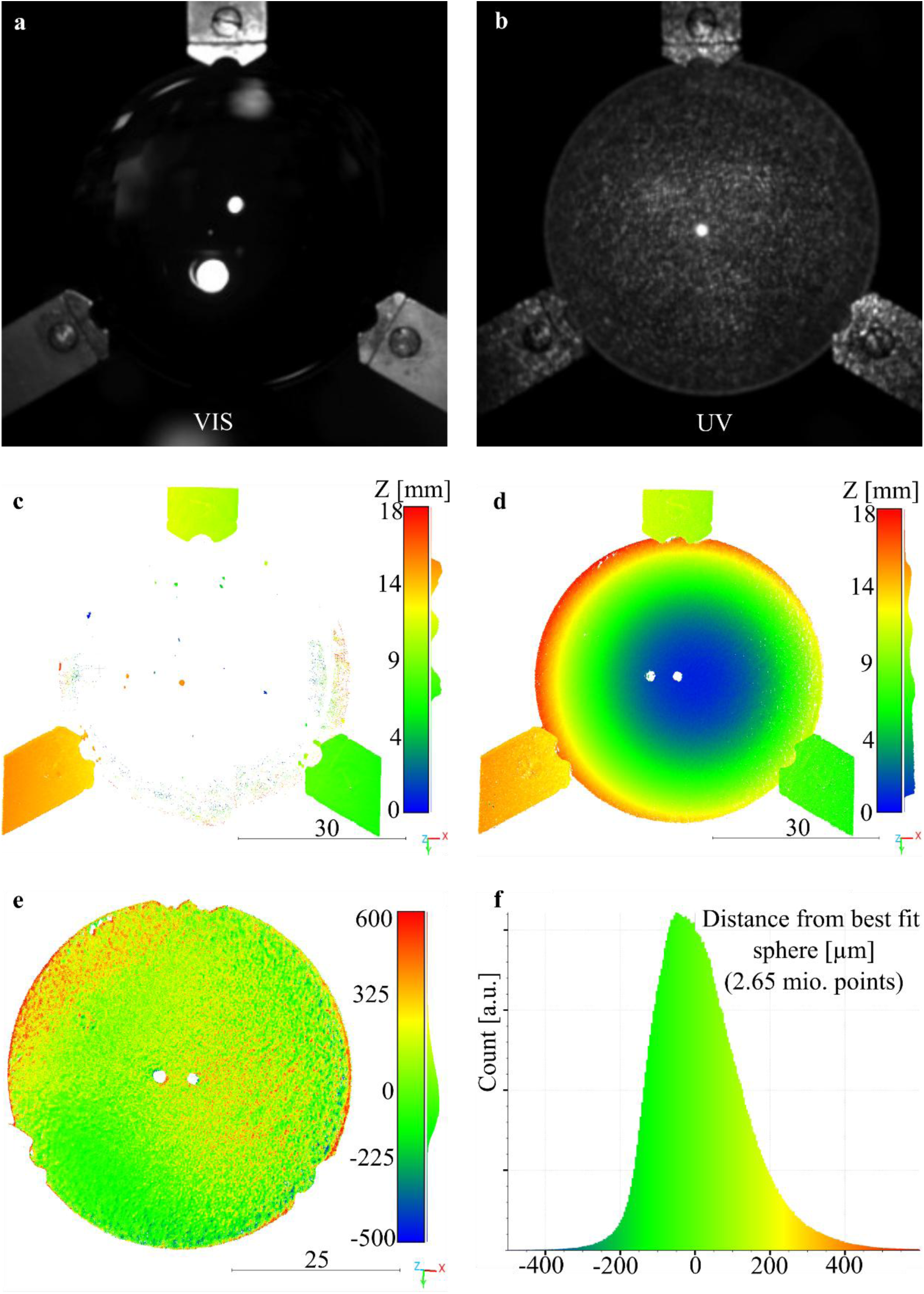
Ultraviolet (UV) speckle illumination enables 3D reconstruction of a transparent glass lens. Comparison of stereo photogrammetric reconstruction of an uncoated plano-convex glass lens (focal length 60 mm, diameter 2-inch, curvature radius ≈ 30.9 mm) using visible (VIS) structured illumination and UV speckle illumination at 266 nm. **(a,b)** Representative camera images under VIS structured illumination (a) and UV speckle illumination (b). While the lens surface appears largely featureless under VIS illumination, UV illumination produces spatial intensity variations due to wavelength-dependent scattering. Exemplary images are available under^20^. **(c,d)** Reconstructed 3D point clouds colored by axial coordinate *z*. Under VIS illumination (c), only sparse points associated with the holder and background are reconstructed, whereas UV illumination (d) enables dense reconstruction of the spherical lens surface. Missing regions near the center arise from specular reflections observed in both camera views. **(e)** Deviation of the UV-based reconstruction from a best-fit sphere. The fitted radius is 30.946 mm with a standard deviation of 106.5 µm, demonstrating accurate recovery of the lens geometry. **(f)** Histogram of point-wise deviations from the best-fit sphere, indicating a narrow distribution centered close to zero. The point clouds have been processed with CloudCompare (www.cloudcompare.org) and are available in the supplementary file^21^.

Beyond enabling reconstruction, UV illumination also allows the detection of small-scale surface defects on transparent materials. This is demonstrated in Fig. 2 for a standard microscopy cover glass.

**Figure 2.**
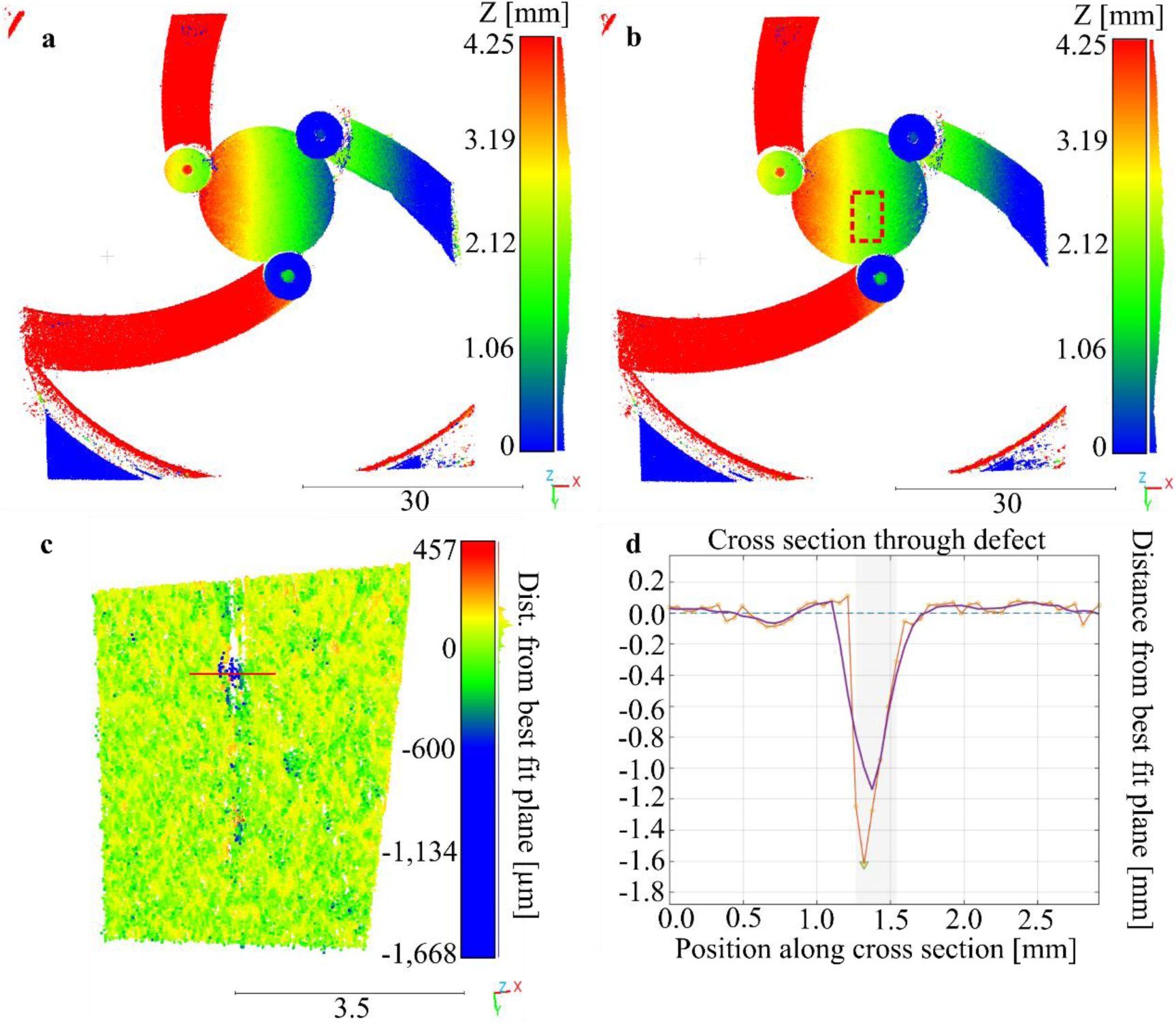
Reconstruction of a transparent cover glass and detection of surface damage using UV speckle illumination. **(a)** Reconstruction of the undamaged cover glass and three-point holder, colored by axial coordinate *z*. The surface appears continuous and homogeneous, with only minor deviations from planarity. The undamaged sample exhibits a standard deviation of 79 µm with respect to a best-fit plane. **(b)** Reconstruction of the same cover glass in the same holder after introducing a surface defect. A vertical cut is clearly visible, slightly below the center of the sample. The area with the defect is marked with a dashed red rectangle. **(c)** Cut-out of the reconstruction of the damaged cover glass as implied by the dashed rectangle in (b), colored by axial coordinate *z*. A red line depicts the points that are shown in the cross section. (**d**) Cross section of the points along the red line from (c), demonstrating that UV illumination enables not only the reconstruction of transparent surfaces but also the detection of small-scale surface defects. The orange line connects the 3D-point positions. The purple line indicates smoothed distribution while the dashed blue line represents the position of the best fit plane. The deepest part of the defect was reconstructed with a distance to the best fit plane of 1.6 mm. Obviously this exceeds the thickness of the cover glass largely, meaning that the defect has been identified correctly but the form of the defect has not been measured accurately.

The undamaged sample is reconstructed as a continuous planar surface with a standard deviation of 79 µm relative to a best-fit plane (Fig. 2a), confirming stable and low-noise reconstruction. After introducing a defect, the same measurement procedure reveals a clearly visible vertical cut in the reconstructed geometry (Fig. 2b–d). Fig. 2 d shows a cross section through the reconstructed points close to and on the defect. Some of the points within the defect are reconstructed outside the actual physical position of the cover glass, that had a thickness of 0.175 mm.

Notably, the defect is resolved without any surface preparation or coating, and despite the inherently low scattering properties of the glass. This highlights that UV speckle illumination not only enables reconstruction of otherwise inaccessible surfaces but also provides sufficient sensitivity for defect detection in transparent components.

To evaluate the applicability of the method to biological specimens with mixed optical properties, a semi-translucent butterfly (*Oleria athalina banjana*, Lepidoptera: Nymphalidae, Ithomiinae) was measured under both coherent VIS and UV speckle illumination (Fig. 3). The result is instructive in a direction not initially anticipated: both modalities successfully reconstruct the specimen, including regions of partial wing transparency, and the cloud-to-cloud distance analysis (Fig. 3d) shows that most surface regions agree closely between VIS and UV reconstructions. Systematic deviations are confined to areas of wing venation, where UV illumination reveals finer structural detail, likely due to enhanced scattering at shorter wavelengths.

**Figure 3.**
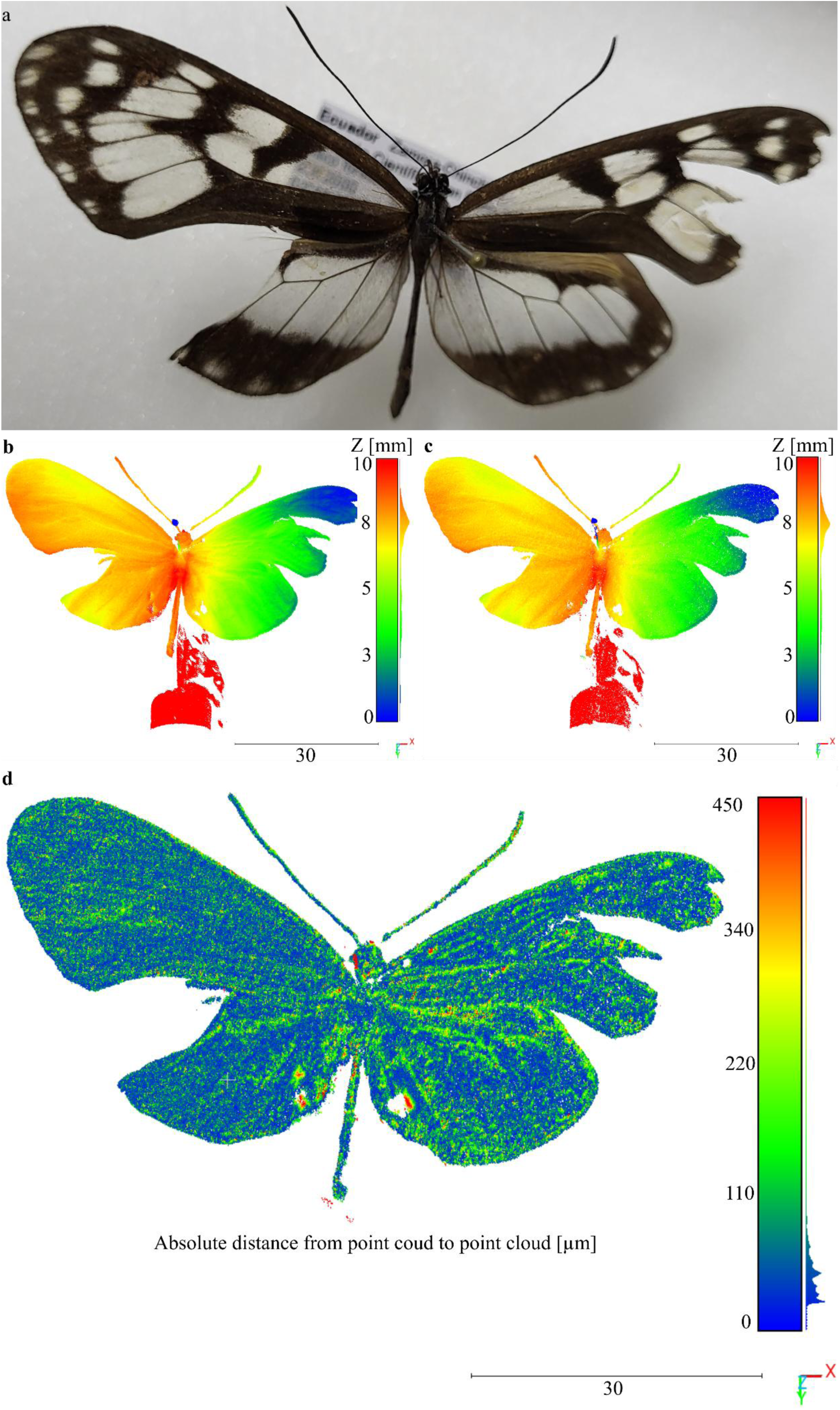
Comparison of 3D-reconstructions of biological composite surfaces in VIS and UV. Photograph **(a)** and stereo photogrammetric reconstruction of a butterfly specimen (*Oleria athalina banjana*, Lepidoptera: Nymphalidae, Ithomiinae) under VIS and UV illumination. **(a)** Photograph of the butterfly specimen. Some parts of the wings are partially transparent. Also the wing venation structures are clearly visible. It is noteworthy that the specimen under test was slightly damaged on the right wing. **(b,c)** Reconstructed 3D point clouds under coherent **VIS** illumination **(b)** and UV speckle illumination at 266 nm **(c)**, colored by axial coordinate z. **(d)** Cloud-to-cloud (C2C) absolute distance between VIS- and UV-based reconstructions. While most regions agree closely, systematic deviations are observed in areas corresponding to wing venation structures. These results indicate that UV illumination reveals additional structural information in biological specimens, likely due to wavelength-dependent scattering and absorption effects. Across all investigated scenarios, UV speckle illumination consistently enables reliable stereo reconstruction in situations where VIS illumination fails.

The fact that both modalities perform comparably on this specimen, in contrast to the complete VIS failure observed on glass surfaces, reflects an important mechanistic distinction. The reconstruction success at 266 nm on the butterfly is a geometrically and physically driven effect: the wing membrane, while partially transparent, provides sufficient scattering at both wavelengths to support correspondence estimation. This is not a spectral-biology effect. Notably, 266 nm lies outside the biologically relevant UV window for UV-capable animals such as insects and birds, which exhibit distinct optical features in the 300 - 400 nm range. The present result therefore establishes a physical baseline: UV photogrammetry at 266 nm reconstructs biological composite surfaces robustly, independently of species-specific UV absorption or reflectance features. This baseline is a prerequisite for future experiments exploiting wavelength-dependent biological contrast, as discussed below.

### Reconstruction accuracy on metrology-grade reference objects

To establish a quantitative performance baseline independent of object-specific effects, metrology-grade reference objects, a plane (surface roughness < 1 µm) and a sphere (radius 15 mm, roughness < 3 µm), were measured under all three illumination modalities under identical geometric and acquisition conditions. These objects provide known ground-truth geometry against which reconstruction accuracy can be assessed as standard deviation from a best-fit reference surface, capturing the combined effect of calibration uncertainty, speckle-induced noise, and correspondence estimation error across the full field of view. The results establish the operating envelope of the system under cooperative scattering conditions and contextualise the accuracy achieved in the transparent and composite object demonstrations above.

The achievable accuracy in practice is influenced by calibration uncertainties, feature size of the projected patterns, image noise, speckle decorrelation, surface scattering behaviour, and correspondence estimation errors. From a purely geometric perspective, the depth sensitivity of the system can be estimated using the stereo triangulation model. Based on the given parameters (focal length 25 mm, pixel pitch 2.74 µm, baseline 95.8 mm, working distance 250 mm), a sensitivity of approximately 71 µm per pixel disparity is obtained. Assuming subpixel correspondence estimation on the order of 0.1 px, this corresponds to an idealized depth resolution of approximately 7 µm.

However, this value represents a local best-case estimate for individual point correspondences under ideal conditions. For practical 3D measurements, particularly over extended fields of view, additional error sources accumulate and introduce both random and systematic deviations. Therefore, the standard deviation of reconstructed points with respect to a known reference geometry (e.g., a best-fit plane or sphere) provides a more representative measure of the achievable precision. This metric captures the combined effect of all error contributions and thus offers a more realistic assessment of the performance of the 3D measurement system in practical scenarios.

To experimentally assess the reconstruction performance under controlled conditions, metrology-grade reference objects, a plane (surface roughness < 1 µm) and a sphere (radius 15 mm, roughness < 3 µm), were measured under identical geometric conditions using all three illumination modalities. For each configuration, 100 stereo image pairs were recorded and processed using the same reconstruction pipeline.

Figures 4 and 5 show representative input images, reconstructed surfaces, and corresponding deviation histograms for the plane and sphere, respectively. VIS structured illumination provides high contrast on the scattering reference objects and results in the most complete and stable reconstructions. In contrast, coherent speckle illumination at 532 nm and 266 nm exhibits reduced completeness and increased noise, which can be partially mitigated by spatial filtering of the input images.

**Figure 4.**
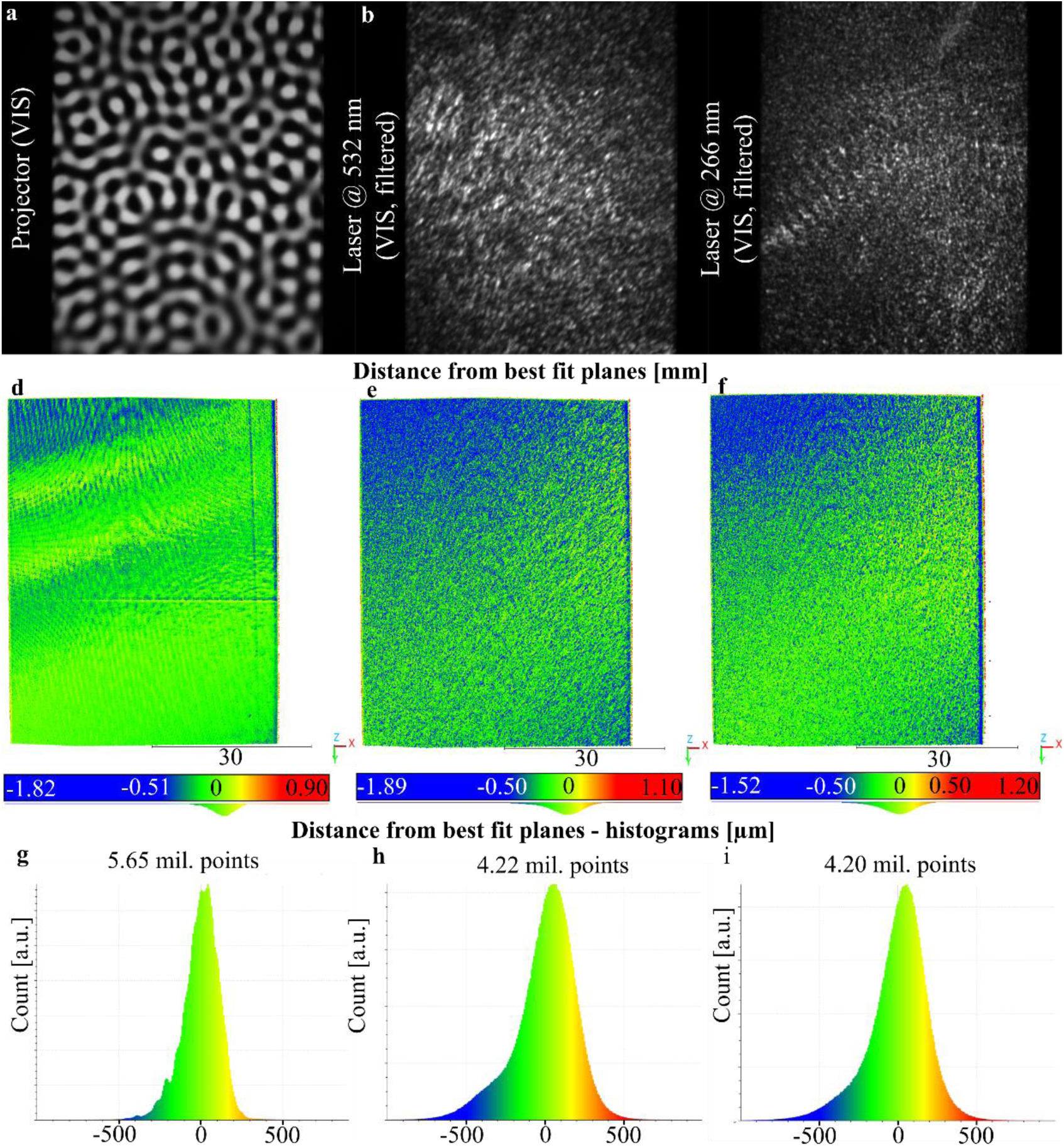
Appearance and reconstructions of metrology-grade reference plane under different illumination modalities. Representative camera images of a metrology-grade plane (roughness < 1 µm) under three illumination conditions. **(a–c)** Plane under broadband VIS structured illumination **(a)**, coherent speckle illumination at 532 nm **(b)**, and at 266 nm **(c)**. Coherent speckle images were spatially filtered using a 1+cos filter (kernel size 21 pixels). **(d–f)** Corresponding images of the 3D-reconstructions of the plane. While VIS structured illumination provides high contrast on scattering surfaces, UV speckle illumination produces comparable contrast with different spatial statistics, enabling robust correspondence estimation. **(g–i)** Histograms corresponding to the reconstructions done with illumination of the incoherent projector in VIS, the speckles at 532 nm and the speckles at 266 nm. The reconstruction results in terms of standard deviation and completeness are shown in Tab. 1.

**Figure 5.**
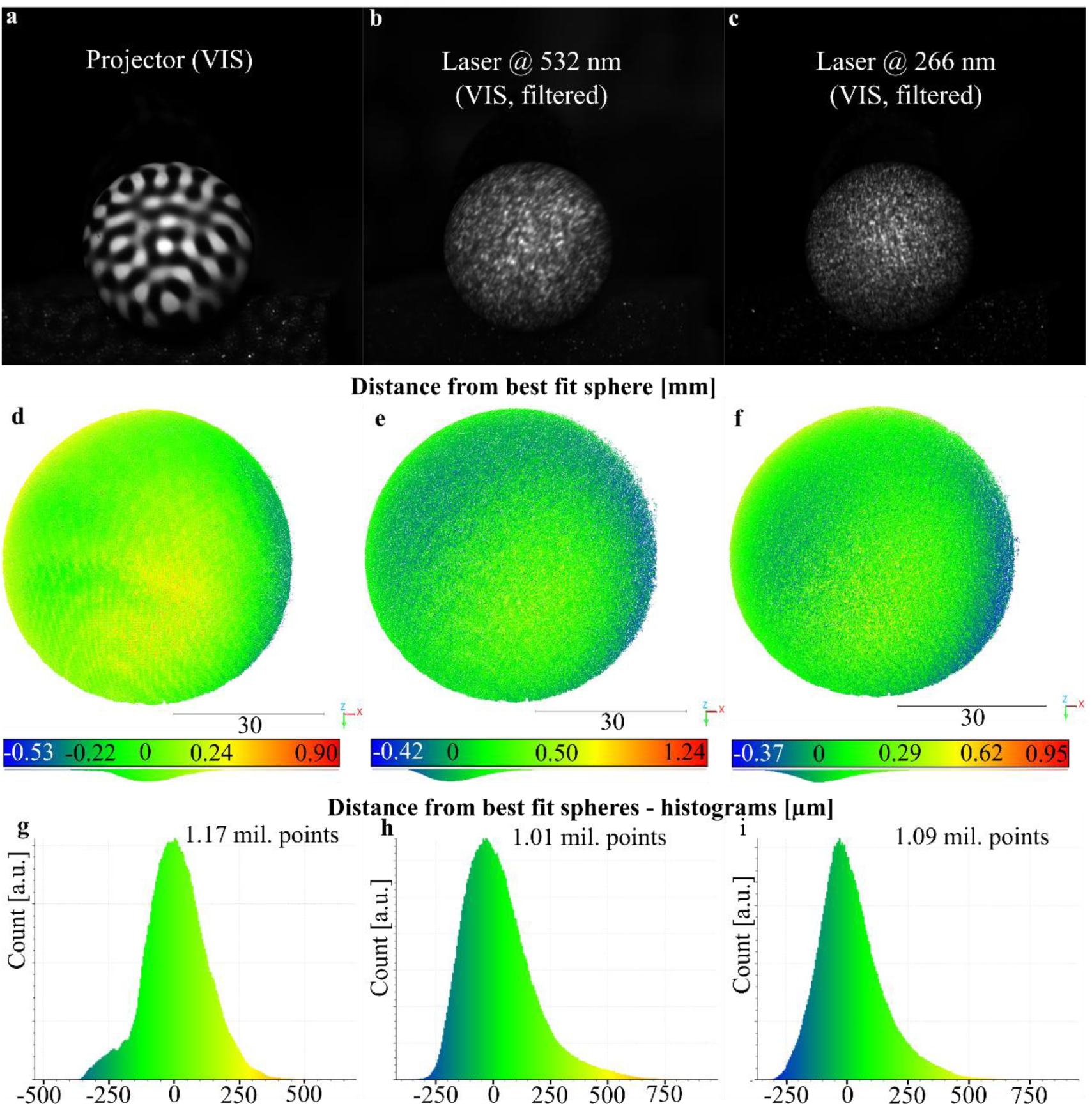
Appearance and reconstructions of metrology-grade reference sphere under different illumination modalities. Representative camera images of a metrology-grade sphere (radius 15 mm, roughness < 3 µm) under three illumination conditions. (a–c) Sphere under broadband VIS structured illumination (a), coherent speckle illumination at 532 nm (b), and at 266 nm (c). Coherent speckle images were spatially filtered using a 1+cos filter (kernel size 21 pixels). (d–f) Corresponding images of the 3D-reconstructions of the sphere. (g–i) Histograms corresponding to the reconstructions done with illumination of the incoherent projector in VIS, the speckles at 532 nm and the speckles at 266 nm. The reconstruction results in terms of standard deviation and completeness are shown in Tab. 1.

**Table 1:**
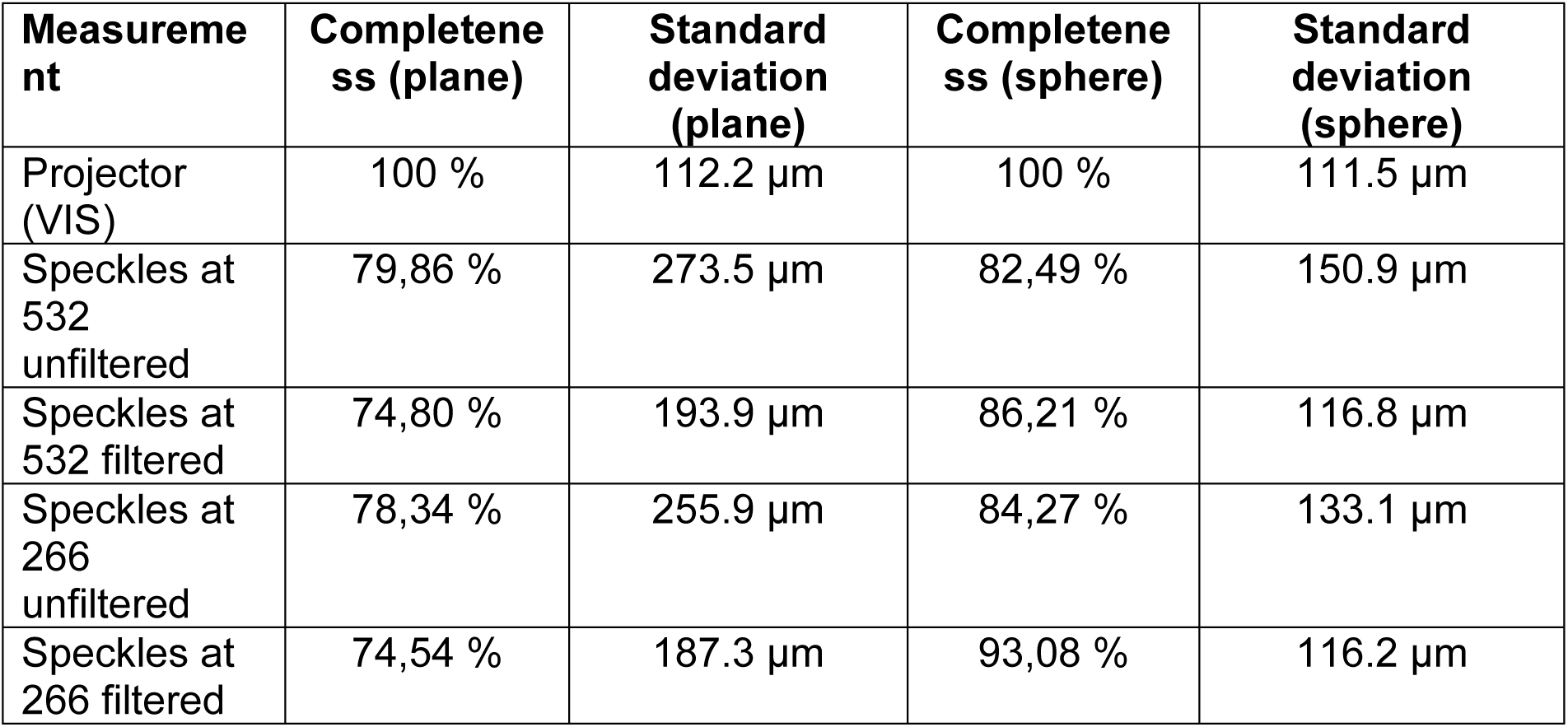
Comparison of 3D-point clouds of metrology grade plane and sphere, taken from 100 image pairs with unchanged geometry. An incoherent illumination with band-limited patterns, a coherent illumination with speckles at 532 nm and another coherent illumination with speckles at 266 nm has been applied. Please note that the terms “unfiltered” and “filtered” are subject to the filtering of the original images, not the point clouds.

The quantitative results summarized in Table 1 confirm these observations. For the plane, VIS structured illumination achieves full completeness (100%) and a standard deviation of 112 µm. Coherent speckle illumination at 532 nm yields lower completeness (≈80%) and higher deviations, which improve after filtering but remain above the performance of VIS structured illumination. UV speckle illumination at 266 nm shows comparable behaviour, with slightly improved performance after filtering (standard deviation ≈187 µm). These comparatively higher deviations for planar surfaces are consistent with the distinction between local depth sensitivity and global surface precision discussed above. While the theoretical depth resolution reflects the sensitivity of individual point correspondences, the reconstruction of extended planar surfaces accumulates residual errors from calibration, correspondence estimation, and speckle-induced intensity variations across the full field of view. In addition, planar geometries are particularly sensitive to low-frequency systematic errors, which directly translate into increased standard deviation with respect to a best-fit reference. Consequently, the observed deviations should be interpreted as a realistic measure of system-level performance rather than a limitation of the underlying depth sensitivity.

For the sphere, a similar trend is observed. VIS structured illumination again achieves full completeness and a standard deviation of approximately 112 µm. Speckle-based approaches show reduced completeness in the unfiltered case but benefit from filtering. Notably, UV speckle illumination achieves a completeness of over 93% after filtering, with a standard deviation of approximately 116 µm, approaching the performance of VIS structured illumination.

Overall, these results indicate that, under cooperative conditions, incoherent VIS structured illumination remains the most accurate approach. However, UV speckle illumination provides comparable reconstruction accuracy while maintaining high completeness, particularly for curved surfaces. Importantly, this performance is achieved under non-optimized acquisition conditions and with identical parameters across all illumination modalities.

These findings support the conclusion that UV photogrammetry offers a competitive level of accuracy while significantly extending the class of measurable objects beyond those accessible with conventional VIS illumination.

## Discussion

The experiments demonstrate three key capabilities:

1. **Reconstruction of transparent surfaces**, including optically smooth glass components, without surface preparation (Fig. 1).
2. **Detection of surface defects** on transparent materials with micrometer-scale sensitivity (Fig. 2).
3. **Improved reconstruction of composite and biological surfaces**, revealing additional structural details (Fig. 3).

In combination with the quantitative evaluation on reference objects (Figs. 4 and 5), these results establish UV photogrammetry as a robust and versatile extension of conventional stereo reconstruction techniques. Taken together, these results indicate that UV speckle illumination fundamentally extends the applicability of stereo photogrammetry beyond its conventional limits, enabling robust reconstruction of transparent and composite surfaces. They also indicate that a combined use of measurement with multiple wavelengths can significantly improve the use-case of stereo photogrammetry.

The presented results demonstrate that UV speckle illumination fundamentally extends the applicability of stereo photogrammetry beyond its conventional limits. While VIS structured illumination fails to provide sufficient contrast on optically smooth transparent surfaces, UV illumination generates measurable back-scattered signals that enable robust correspondence estimation and dense 3D reconstruction. The observed UV-enhancement of backscattered signal is consistent with wavelength-dependent surface scatter theory. For an optically smooth surface with RMS roughness σ_r_, scatter intensity in the Beckmann-Kirchhoff regime scales approximately as (σ_r_/λ)^2^, predicting a factor of four increase in scattered intensity when the illumination wavelength is halved from 532 nm to 266 nm, independent of bulk material properties^18^. For subsurface inhomogeneities and bulk scatter contributions, Rayleigh-type scaling provides an additional enhancement, with scatter intensity following an approximate λ^−4^ dependence^18,19^. Both mechanisms act simultaneously in optical glass, and their combined effect at 266 nm relative to the visible range is sufficient to produce measurable backscatter from surfaces that are effectively non-scattering under VIS illumination. Experimental evidence for UV-enhanced scatter in optical glass at deep UV wavelengths has been reported in the literature: Schröder et al. demonstrated substantial bulk scatter in synthetic fused silica at 193 nm, confirming that UV-induced scattering is a material-level phenomenon in optical glasses rather than an artefact of surface preparation^19^. In the present experiments, the transition from zero detectable signal under 532 nm illumination to dense reconstruction under 266 nm illumination is consistent with crossing a detection threshold governed by this wavelength-dependent scatter enhancement. A full spectral characterization of the backscatter intensity as a function of wavelength for the specific glass materials used, which would allow direct comparison with theoretical predictions, remains an open experimental task and a priority for future work.

Beyond enabling the reconstruction of transparent objects, the experiments show that UV speckle illumination remains compatible with scattering surfaces. This allows the reconstruction of composite surfaces that simultaneously exhibit transparent, reflective, and diffuse regions, scenarios that are challenging for many established optical metrology techniques. Structured light projection requires sufficient surface contrast, deflectometry relies on specular reflections, and interferometric techniques often demand well-controlled optical conditions. In contrast, the presented approach enables the recovery of the overall geometry, position, and defect structure of such composite objects within a single measurement framework, without the need for surface preparation or coating. This capability is particularly relevant for industrial environments, where robust and flexible measurement solutions are required.

The cover glass experiment illustrates an inherent limitation of stereo photogrammetry when applied to thin, highly transparent objects with sharp geometric discontinuities. The reconstructed depth of the defect (1.6 mm)substantially exceeds the physical thickness of the cover glass (0.175 mm), indicating that the correspondence estimator assigns incorrect matches in the vicinity of the defect edge. This arises because the NCC-based matching relies on local intensity patterns: near a sharp crack or edge, the backscattered signal from the defect walls, the opposing surface, and potential multiple-reflection paths can generate misleading texture that leads to erroneous disparity estimates. The result is that the defect is reliably detected and localized in-plane, but its three-dimensional shape and depth cannot be recovered accurately by the present pipeline. This distinction — between detection capability and geometric fidelity — is important for the intended application context: UV photogrammetry is well suited as a screening tool to identify and locate defects on otherwise inaccessible transparent surfaces, while quantitative characterization of defect geometry would require a complementary high-precision technique such as confocal microscopy or interferometry.

From an application perspective, the method is well suited for scenarios where a reliable “coarse” 3D description is sufficient or serves as a prerequisite for further processing. This includes applications in industrial inspection, such as 100% quality control, defect detection on transparent components, and geometry verification in pick-and-place operations. In such contexts, the ability to reconstruct otherwise uncooperative surfaces can close an important gap between vision-based inspection and precision metrology.

At the same time, the approach is inherently compatible with more precise measurement techniques. The presented method can provide an initial geometric estimate or region-of-interest detection that can subsequently be refined using techniques such as deflectometry, interferometry, or infrared-based 3D sensing. In this sense, UV photogrammetry can act as an enabling front-end within multimodal measurement pipelines, combining robustness and flexibility with the precision of specialized techniques.

It is important to note that the presented setup was not optimized for maximum performance of each illumination modality. Instead, identical acquisition parameters, including exposure times, were used across VIS structured illumination, 532 nm speckle illumination, and 266 nm speckle illumination to ensure comparability. This design choice is reflected in the quantitative results on metrology-grade objects, where VIS structured illumination outperforms coherent speckle illumination under cooperative conditions. The results therefore represent a conservative comparison rather than the achievable performance limits of each modality.

In conclusion, UV illumination allows stereo photogrammetry to reconstruct transparent and composite surfaces that are not accessible under conventional VIS illumination. The presented approach enables robust reconstruction of geometry and defects without surface preparation while maintaining competitive accuracy under non-optimized conditions. As a complementary technique, UV photogrammetry can serve as a flexible front-end for industrial inspection tasks and as an entry point for multimodal measurement pipelines combining robustness with high-precision methods. These results highlight the potential of wavelength-adapted illumination strategies to broaden the scope of optical 3D metrology across industrial and biological applications. Further improvements can be expected from optimized illumination concepts. In particular, the use of incoherent or broadband UV illumination may reduce speckle-induced noise^22,23^ and improve reconstruction accuracy, analogous to the advantages of incoherent structured illumination in the VIS range. The development of such illumination systems remains a technical challenge but represents a promising direction for future work.

Beyond industrial applications, the biological measurements presented here open a specific direction for future work. The current experiments at 266 nm establish that UV photogrammetry reconstructs biological composite surfaces through a geometry-driven scattering mechanism, without dependence on species-specific spectral features. This is a necessary and non-trivial baseline: it demonstrates that the technique is robust across surface types before wavelength-selective biological contrast is introduced. The next logical step is to shift illumination into the near-UV range, approximately 300 - 400 nm, where many organisms, particularly insects and flowering plants, exhibit pronounced wavelength-dependent optical properties including differential UV reflectance, structural coloration, and UV-absorbing pigmentation^24–27^. At these wavelengths, transparent and absorbing wing regions that are indistinguishable geometrically may produce differential scattering signatures, potentially enabling functional as well as geometric reconstruction within a single measurement. A wavelength-tunable incoherent UV source in this range would combine the speckle-noise advantages of incoherent illumination with spectral selectivity and represents the most direct technical extension of the present system.

## Materials and Methods

### Experimental setup

The experimental setup consists of a stereo photogrammetric system with three illumination pathways. The details have been listed in Tab. 2.

**Table 2:**
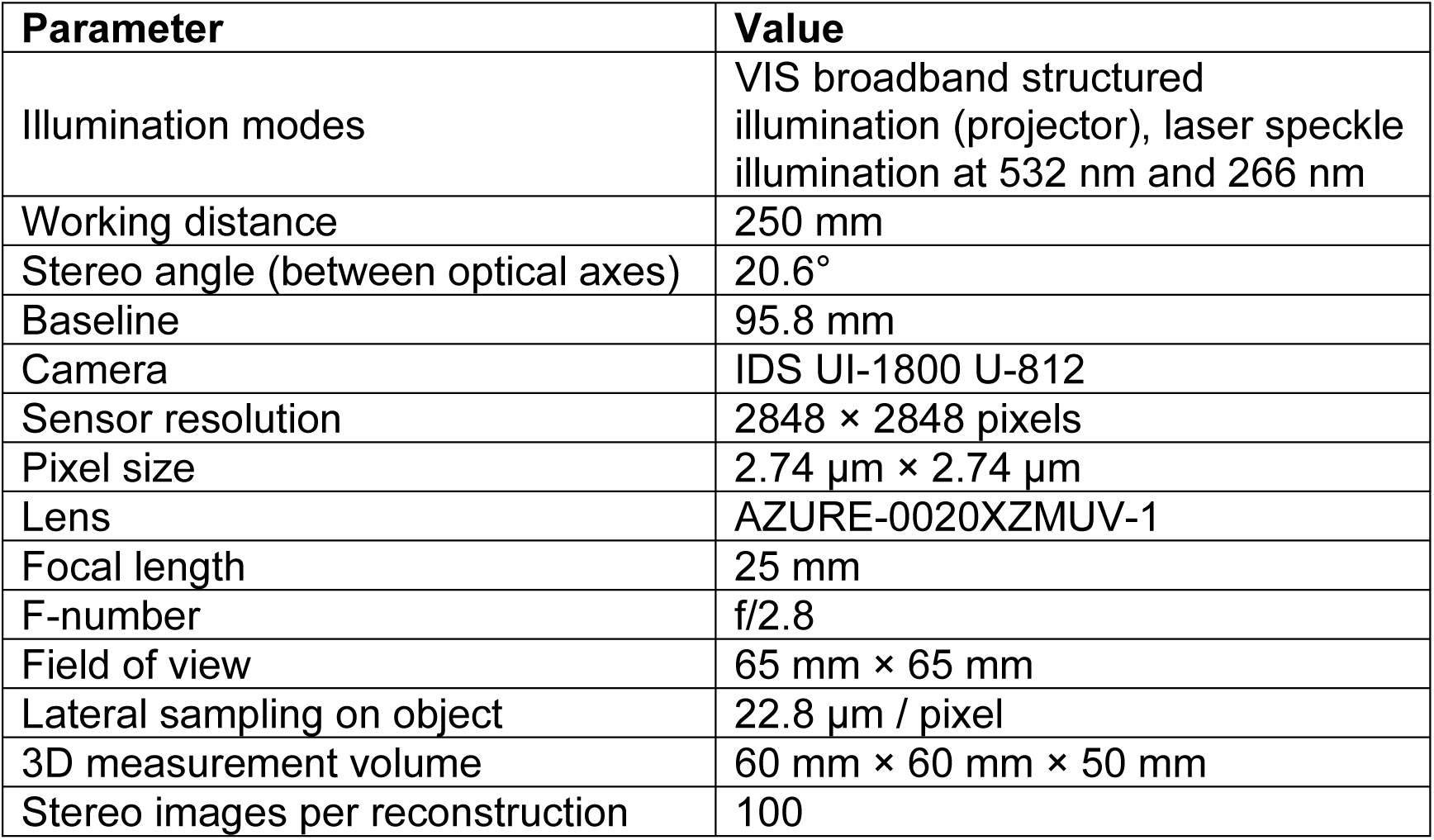
Specifications of the measurement setup.

| Parameter | Value |
| --- | --- |
| Illumination modes | VIS broadband structured illumination (projector), laser speckle illumination at 532 nm and 266 nm |
| Working distance | 250 mm |
| Stereo angle (between optical axes) | 20.6° |
| Baseline | 95.8 mm |
| Camera | IDS UI-1800 U-812 |
| Sensor resolution | 2848 × 2848 pixels |
| Pixel size | 2.74 μm × 2.74 μm |
| Lens | AZURE-0020XZMUV-1 |
| Focal length | 25 mm |
| F-number | f/2.8 |
| Field of view | 65 mm × 65 mm |
| Lateral sampling on object | 22.8 μm / pixel |
| 3D measurement volume | 60 mm × 60 mm × 50 mm |
| Stereo images per reconstruction | 100 |

Three illumination modes were implemented in order to investigate different surface scattering conditions: broadband VIS structured illumination generated by an of-the-shelf projector (Technaxx Mini LED Beamer TX-113) and an additional lens, laser speckle illumination at 532 nm (MGL-F-532-2W), and UV speckle illumination at 266 nm (MPL-N-266). Structured illumination sequences consisting of 100 patterns were recorded for each measurement. The acquisition time for each illumination was fixed independent of the object. For the incoherent projector an acquisition time of 17 ms and for the speckle illumination (at 532 nm and at 266 nm) an acquisition time of 450 ms have been used for each image. The speckle patterns have been changed by using a wobble-mirror. With a rotational motor the mirror was brought into different orientation therefore changing the light path of the laser far enough to change the speckle patterns, while moving the laser the image acquisition was halted. The UV-Laser has been used at maximum intensity of 100 mW while the 532nm-Laser (MGL-F-532-2W) has been tuned down to match the average intensity when illuminating the reference plane. Fig. 6. shows a sketch of the setup in use.

**Figure 6.**
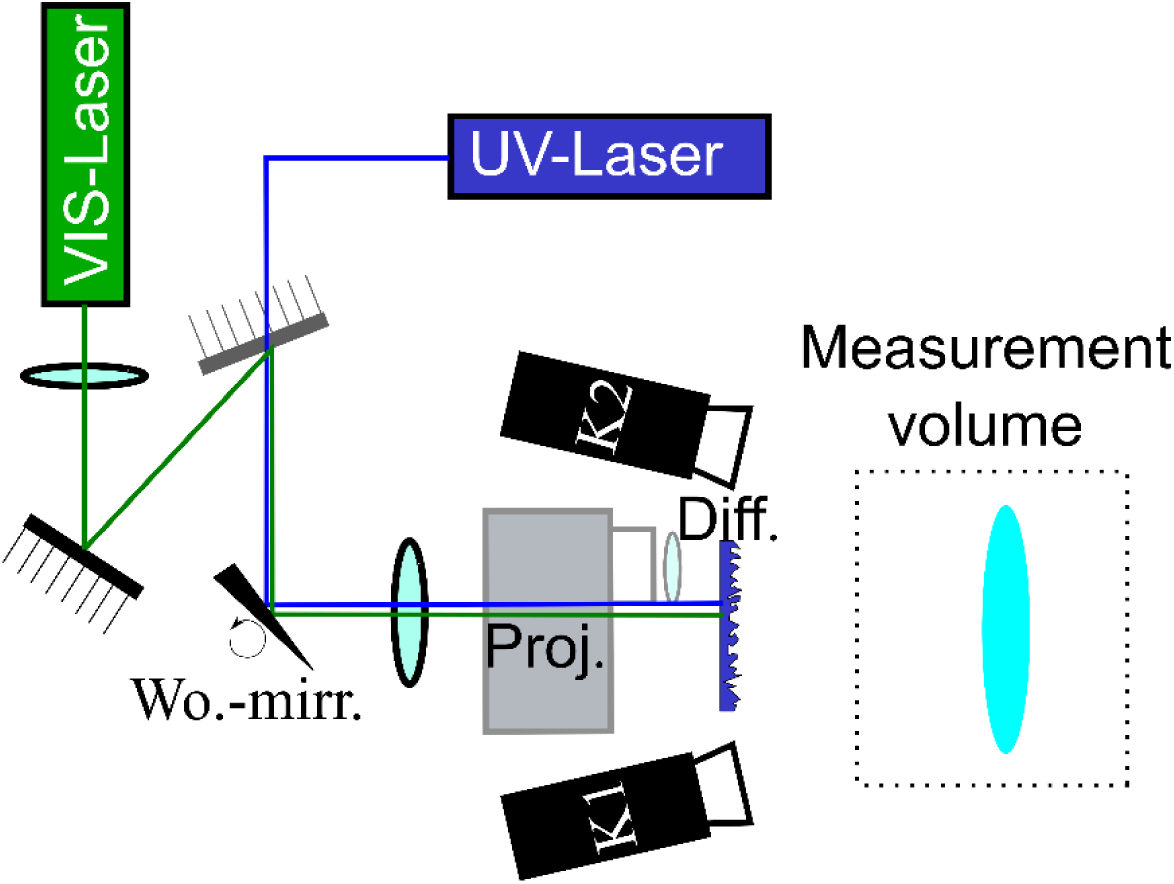
Experimental setup for multispectral stereo photogrammetry. Schematic illustration of the measurement system. Two cameras (K1, K2), sensitive in both VIS and UV spectral ranges, are arranged in a stereo configuration. Illumination is provided either by a broadband VIS projector (Proj.) or by coherent laser sources at 532 nm (VIS) and 266 nm (UV). Speckle patterns are generated using a diffuser (Diff.) and dynamically varied by a wobble mirror (Wo.-mirr.), which changes the optical path of the laser beam. The measurement volume is indicated by the dashed box. The cameras simultaneously observe the illuminated object, and 3D reconstruction is performed using correlation-based correspondence estimation and stereo triangulation. Please note that the projector was positioned above the projection path of the laser speckles to avoid obstruction.

### Filtering of the speckle image

To improve the robustness of correspondence estimation under coherent illumination, the recorded speckle images were subjected to a two-stage filtering procedure.

First, a spatial smoothing filter was applied to reduce high-frequency speckle noise while preserving larger-scale intensity variations relevant for correlation. This was implemented as a one-dimensional cosine-based kernel of the form

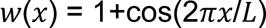

applied in vertical direction with a kernel size of 21 pixels^28^. The filter acts as a band-limited low-pass operator, attenuating fine-scale speckle fluctuations while maintaining spatial structures that contribute to stable matching. Second, a temporal filtering step was performed across the image stack acquired for each measurement, also for the incoherent measurements. For each pixel, the standard deviation over all recorded frames was computed, yielding a statistical measure of intensity variation induced by the changing speckle patterns. Static background contributions and low-variance regions were suppressed by weighting or thresholding based on this per-pixel standard deviation, thereby enhancing regions with significant temporal modulation. This combined spatial–temporal filtering approach reduces speckle-induced decorrelation effects and background noise, resulting in improved stability and accuracy of the subsequent normalized cross-correlation (NCC)-based correspondence estimation.

Quantitative reconstruction accuracy was evaluated on metrology-grade reference objects; results are reported in the Results section.

### Reconstruction pipeline

The 3D-reconstruction is performed using a correlation-based stereo photogrammetry pipeline consisting of four main stages: image acquisition and rectification, correspondence estimation, triangulation, and point cloud generation.

In the first step, synchronized stereo image sequences are recorded for each illumination modality. For structured illumination and speckle-based measurements, multiple images are acquired while varying the projected pattern to enhance feature diversity and improve matching robustness. The images are rectified using the same intrinsic and extrinsic parameters for all illumination types. This is possible because the objectives used are highly corrected across the UV-to-VIS spectral range.

In the second step, correspondences between the two camera views are established using normalized cross-correlation (NCC). The matching is performed on temporal intensity patterns, depending on the illumination modality. For each pixel in one camera image, a corresponding location in the second image is determined by maximizing the NCC score within a predefined search region.

Based on the established correspondences, 3D points are computed by stereo triangulation using the calibrated camera parameters. The reconstruction is performed in the original camera coordinate system, ensuring geometric consistency with the calibrated setup.

Finally, the resulting 3D points are aggregated into a dense point cloud. Each point is associated with its spatial coordinates, correlation score, and optionally intensity information from the input images. The resulting point cloud forms the basis for subsequent analysis, such as surface fitting, deviation mapping, and defect detection.

### Limitations and optimization strategies for the current setup

The presented setup was not optimized for maximum performance of the individual illumination modalities. Instead, identical acquisition parameters, including illumination intensity and exposure time (this separate for the incoherent case –17 ms- and the coherent cases −450 ms), were deliberately used across all projection types to ensure a consistent and fair comparison. As a result, the system was optimized with respect to comparability rather than absolute performance.

Under these conditions, incoherent VIS structured illumination outperforms coherent speckle-based approaches on cooperative scattering surfaces, as expected. The coherent illumination schemes at 532 nm and 266 nm exhibit comparable behaviour, with performance primarily limited by speckle-induced noise and decorrelation effects. These observations indicate that the reported results represent a conservative estimate of the achievable performance, particularly for UV illumination.

Further improvements can be expected from optimized illumination strategies.

In particular, the implementation of an incoherent or broadband UV illumination system could reduce speckle-related artefacts and improve reconstruction accuracy, analogous to the advantages of incoherent structured illumination in the VIS spectral range. Such a system could additionally incorporate multiple wavelengths within the UV and near-UV regime.

This is of particular interest for biological applications, as many organisms, especially insects and birds, exhibit pronounced wavelength-dependent optical properties in the spectral range between 300 nm and 400 nm. A wavelength-tunable UV illumination system may therefore provide additional contrast mechanisms and enable enhanced structural analysis.

More generally, UV stereo photogrammetry should be regarded as a complementary technique that extends, rather than replaces, existing optical metrology approaches. It can be combined with VIS and infrared imaging as well as with high-precision methods such as deflectometry or interferometry. In this context, UV photogrammetry can provide robust initial geometry, defect localization, or region-of-interest detection, which can subsequently be refined using more specialized measurement techniques.

In the long term, this integration of robustness and flexibility may enable new applications in industrial environments, including automated inspection, 100% quality control, and geometry verification in pick-and-place processes, as well as in biological surface analysis.

## Acknowledgements

This work has been supported by German Federal Ministry of Research, Technology and Space (BMFTR) within the Research Program Quantum Systems, program ‘3DVens’ under the project number 13N16890. People who helped with the preparation of the UVLAS-project early to set this research in motion: Dr. Jan Rothhardt for allowing a first test with a laser at 350 nm already in 2016, Dr. Ann-Sophie Munser from making a spectral straylight distribution of certain materials, such as stainless steel and Dr. André Boden, Dr. Daniel Richter and Dr. Maximilian Heck for letting us use a femtosecond laser below 300 nm.

## Author Contributions

C.F. and A.W.S. supervised the whole project. C.F. and A.W.S. conceived the experiments. G.J.G. and A.P. constructed experimental setups. G.J.G. wrote the software to take images and control the projector and the wobble mirror.

A.W.S. carried out experiments. M.G. wrote the reconstruction code. D. B. and G.B. provided insight into biological specimens and UV-measurements thereof. L.H. performed literature review and gave advice on biological interpretation in the context of 3D-data. All authors participated in the analysis of data and contributed to the writing of the manuscript.

## Data availability

Supplementary materials are available at the online version. The 2D-images, the intrinsic and extrinsic parameters, and a MATLAB code for image filtering are available under^20^. The shown 3D-reconstructions are available here^21^.

## References

1. Zhang, S. High-speed 3D imaging with digital fringe projection techniques. In Tribute to James C. Wyant: The extraordinaire in optical metrology and optics education, Proc. SPIE 11813 (2021). 10.1117/12.2567675

2. Heist, S. et al. High-speed three-dimensional shape measurement using GOBO projection. Opt. Lasers Eng. 87, 90–96 (2016). 10.1016/j.optlaseng.2016.02.017

3. Schaffer, M., Grosse, M. & Kowarschik, R. High-speed pattern projection for three-dimensional shape measurement using laser speckles. Appl. Opt. 49, 3622–3629 (2010). 10.1364/AO.49.003622

4. Schaffer, M. et al. High-speed three-dimensional shape measurements of objects with laser speckles and acousto-optical deflection. Opt. Lett. 36, 3097–3099 (2011). 10.1364/OL.36.003097

5. Stark, A. W. et al. Miniaturization of a coherent monocular structured illumination system for future combination with digital holography. Light Adv. Manuf. 3, 437–444 (2022). 10.37188/lam.2022.034

6. Stark, A. W. et al. Telecentric stereo 3D imaging with isotropic micrometer resolution bridges macro- and microscale in small Lepidopterans. Sci. Rep. 15, 28690 (2025). 10.1038/s41598-025-13795-6

7. Weiler, S. et al. A primary sensory cortical interareal feedforward inhibitory circuit for tacto-visual integration. Nat. Commun. 15, 3081 (2024). 10.1038/s41467-024-47459-2

8. Quan, H., Shi, W. & Kong, L. Non-destructive optical measurement of transparent objects: a review. Light Adv. Manuf. 6, 333–357 (2025). 10.37188/lam.2025.022

9. Wu, Z. et al. Dynamic 3D shape reconstruction under complex reflection and transmission conditions using multi-scale parallel single-pixel imaging. Light Adv. Manuf. (2024). 10.37188/lam.2024.034

10. Knauer, M. C., Kaminski, J. & Häusler, G. Phase measuring deflectometry: a new approach to measure specular free-form surfaces. Proc. SPIE 5457 (2004). 10.1117/12.545704

11. Huang, L. et al. Review of phase measuring deflectometry. Opt. Lasers Eng. (2018). 10.1016/j.optlaseng.2018.03.026

12. Lang, W. et al. Deterministic form-position deflectometric measurement of monolithic multi-freeform optical structures via Bayesian multisensor fusion. Light Adv. Manuf. (2025). 10.37188/lam.2025.029

13. Häusler, G. & Willomitzer, F. Reflections about the holographic and non-holographic acquisition of surface topography: where are the limits? Light Adv. Manuf. 3, 226–235 (2022). 10.37188/lam.2022.025

14. de Groot, P. J. et al. Contributions of holography to the advancement of interferometric measurements of surface topography. Light Adv. Manuf. 3, 258–277 (2022). 10.37188/lam.2022.007

15. Fratz, M. et al. Digital holography in production: an overview. Light Adv. Manuf. 2, Article 15 (2021). 10.37188/lam.2021.015

16. Speck, H. et al. Analysis of the measurement accuracy of a thermal 3D sensor for transparent objects. Measurement 258, 119068 (2026). 10.1016/j.measurement.2025.119068

17. Landmann, M. et al. High-resolution sequential thermal fringe projection technique for fast and accurate 3D shape measurement of transparent objects. Appl. Opt. 60, 2362–2371 (2021). 10.1364/AO.419492

18. Bohren, C. F. & Huffman, D. R. Absorption and Scattering of Light by Small Particles. Wiley (1983).

19. Schröder, S. et al. Bulk scattering properties of synthetic fused silica at 193 nm. Opt. Express 14, 10537–10549 (2006). 10.1364/OE.14.010537

20. Gentsch, G. J., et al. Ultraviolet illumination enables 3D-reconstruction of transparent and composite surfaces [Supplementary data]. figshare (2026). 10.6084/m9.figshare.31999497

21. Gentsch, G. J., et al. Ultraviolet illumination enables 3D-reconstruction of transparent and composite surfaces [Data set]. Zenodo (2026). 10.5281/zenodo.19555833

22. Goodman, J. W. Speckle Phenomena in Optics: Theory and Applications, 2nd ed. SPIE Press (2015).

23. Evered, C., Li, K., Fan, Y., Zhang, B. & Roula, A. A review of light sources used for laser speckle reduction in display and imaging applications. Opt. Laser Technol. 183, 112407 (2025). 10.1016/j.optlastec.2024.112407

24. Lind, O., Mitkus, M., Olsson, P. & Kelber, A. Ultraviolet vision in birds: the importance of transparent eye media. Proc. R. Soc. B 281, 20132209 (2014). 10.1098/rspb.2013.2209

25. Briscoe, A. D. & Chittka, L. The evolution of color vision in insects. Annu. Rev. Entomol. 46, 471–510 (2001). 10.1146/annurev.ento.46.1.471

26. Cronin, T. W. et al. Photoreception and vision in the ultraviolet. J. Exp. Biol. 219, 2790–2801 (2016). 10.1242/jeb.128769

27. Pinna, C. S. et al. Mimicry can drive convergence in structural and light transmission features of transparent wings in Lepidoptera. eLife 10, e69080 (2021). 10.7554/eLife.69080

28. Stark, A. W. et al. Subjective speckle suppression for 3D measurement using one-dimensional numerical filtering. Appl. Opt. 58, 9473–9483 (2019). 10.1364/AO.58.009473

